# *Prochlorococcus* predation by a globally abundant filter feeder

**DOI:** 10.1101/2024.11.25.625256

**Authors:** Carey P. Sweeney, Avery Harman, Anvita U. Kerkar, Anne E. Aasjord, Daniel M. Chouroutt, Terra Hiebert, Kelly Sutherland, Anne W. Thompson

## Abstract

*Prochlorococcus* is the most abundant photosynthetic cell on Earth and is critical to primary productivity and biogeochemical cycles of the open ocean. Appendicularians are ubiquitous gelatinous filter-feeding zooplankton that feed on marine microorganisms including *Prochlorococcus*. However, the details of this feeding interaction are extremely understudied relative to its potential importance in top-down controls on *Prochlorococcus*. This is the first study to experimentally examine several dimensions of the feeding interaction between cultivated appendicularians and *Prochlorococcus*. We found that *Prochlorococcus* retention rates by the appendicularian *Oikopleura dioica* increased with prey concentration and predator age. We found that appendicularians grazed equally on the two most globally abundant *Prochlorococcus* ecotypes HLI and HLII and that the presence of larger diatom prey did not change *Prochlorococcus* retention rates. The quantitative insight and retention rates provided by this study will help fill gaps in models of the marine carbon cycle and marine microbial community dynamics and biogeography, and expand the knowledge of *Prochlorococcus* ecology.

**Importance:** Appendicularians are a known predator of picocyanobacteria, but the details of their feeding on the globally abundant picocyanobacterium *Prochlorococcus* have not been investigated. We quantified *Prochlorococcus* retention rates over a range of ecologically relevant conditions, which will inform microbial community predictions and carbon flux models and lead to improved understanding of carbon transfer in the ocean, microbial ecology, and microbial communities.

## Introduction

*Prochlorococcus* is a superabundant microorganism that is forecast to increase in global abundance with climate change (Biller et al., 2015; Flombaum et al., 2013). In vast areas of the open ocean, *Prochlorococcus* is the dominant photosynthetic cell and therefore contributes to primary production on a global scale (Field et al., 1998). Understanding the factors that control *Prochlorococcus* abundance, production, and entry into carbon cycles are key to understanding the ocean system and the ecology of abundant marine microorganisms.

*Prochlorococcus* growth is balanced with its mortality across the open oceans (Ribalet et al., 2015), however the many mechanisms of *Prochlorococcus* mortality are not well-integrated into its ecology. Most mortality of *Prochlorococcus* is attributed to viral lysis (Carlson et al., 2022) and protist predation (Connell et al., 2020), however it is not clear that these mortality sources account for all *Prochlorococcus* death in all areas of the ocean. For example, active viral infections are localized to hot spots (Carlson et al., 2022), and ecosystem models that include viral and protist predation have revealed uncertainties in understanding *Prochlorococcus* mortality (Beckett et al., 2024; Talmy et al., 2019, 2024). Further, mechanisms of cell death influence microbial contributions to the carbon cycle, but this is not well-explored for *Prochlorococcus*. For example, viral lysis produces dissolved organic carbon profiles that are distinct from mechanical cell disintegration (Ma et al., 2018). Thus, examination of novel mortality sources will improve understanding of *Prochlorococcus* ecology, evolution, and contributions to biogeochemical cycles.

Appendicularians are mucous mesh grazers known to remove *Prochlorococcus* from seawater (Dadon-Pilosof et al., 2023; Scheinberg et al., 2005). Cells captured in the appendicularian’s sophisticated filtration system (i.e. mucous house) (Conley et al., 2018; Hiebert et al., 2023) can be digested, rejected (Lombard et al., 2011), incorporated into sinking fecal pellets (Sato et al., 2001), or trapped in discarded houses that become marine snow (Lombard & Kiørboe, 2010) (Figure 1). These discrete pathways for prey particles, after appendicularian capture, lead to unique contributions of appendicularians to global carbon cycles (Jaspers et al., 2023; Taucher et al., 2024). Appendicularians can be highly abundant during blooms (Uye & Ichino, 1995) and in some cases they are the most abundant zooplankton (Schmid et al., 2023). Appendicularian distribution is cosmopolitan (Masunaga et al., 2022) and overlaps with *Prochlorococcus* habitats (Visintini et al., 2021) raising questions about the details and importance of this interaction. Appendicularian feeding is influenced by many factors, which have been revealed through studies on artificial particles, eukaryotic phytoplankton, and large viruses. In laboratory experiments, appendicularian feeding on artificial prey particles is selective based on particle size, shape (Conley & Sutherland, 2017), and surface properties (Hiebert et al., Submitted). Appendicularian feeding also varies with predator characteristics such as age (Troedsson et al., 2009), species (Dadon-Pilosof et al., 2023), and the extent of house clogging by abundant or spiny prey items (Selander & Tiselius, 2003; Troedsson et al., 2007). Understanding how appendicularian feeding dynamics and prey selection apply to marine bacteria, such as *Prochlorococcus,* would bolster understanding of microbial mortality and carbon cycling in the surface ocean for a broad range of marine bacteria.

**Figure 1.**
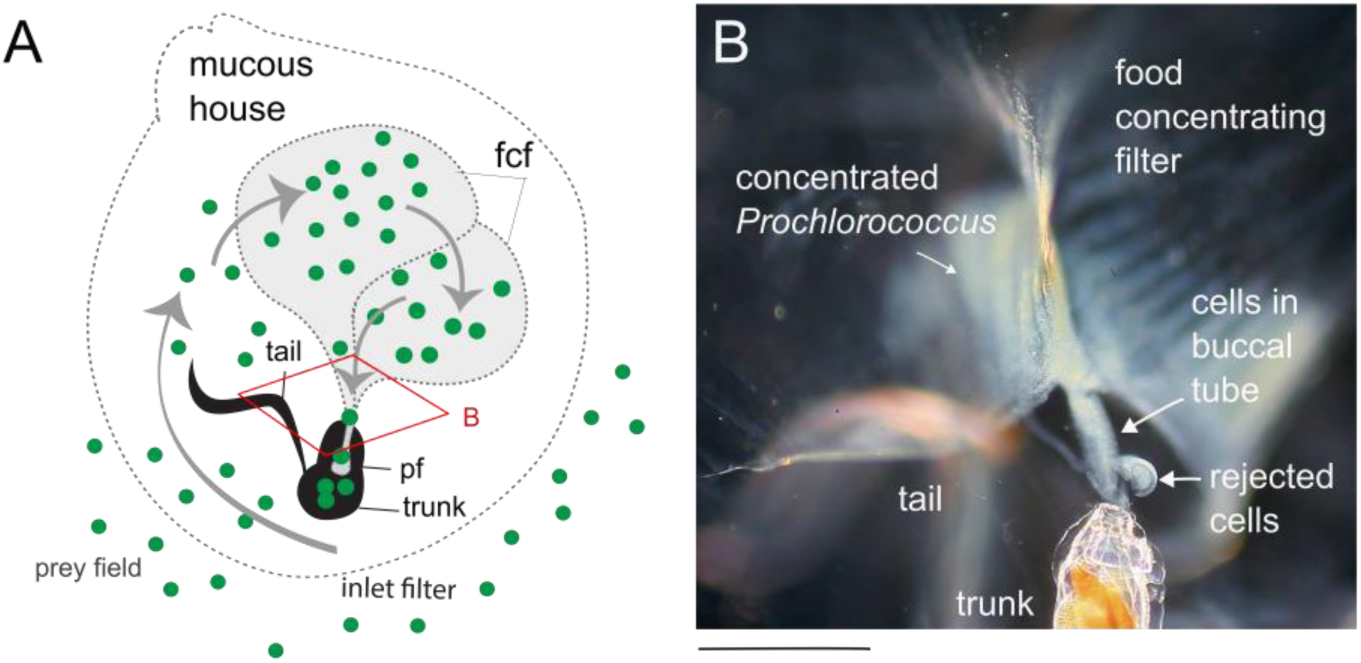
Appendicularian interactions with *Prochlorococcus* from two different views. A) Lateral view of appendicularian feeding and hydrodynamic processes, modified from Hiebert et al. (In review), with *Prochlorococcus* represented as green circles (scaled much larger) and the appendicularian in black (with trunk, inlet filter location, food concentrating filter, pharyngeal filter, and tail labeled). Dashed lines show outlines of select mucous features. Gray arrows show approximate fluid path due to tail beating. Inset (in red), shows orientation angle and area photographed in B. B) Light microscopy (trunk view) of a live appendicularian feeding on *Prochlorococcus* (MIT9312, green-orange appearance due to light artifact). Abbreviations: Food concentrating filter (fcf), pharyngeal filter (pf). Scale bar approximately 500 µm.

This study addresses the details of *Prochlorococcus* mortality by appendicularians. With the cultivated appendicularian *Oikopleura dioica* and cultivated *Prochlorococcus* as prey (Figure 1B), we used qPCR to measure retention rates over a range of ecologically relevant conditions, including ranging prey concentrations, *Prochlorococcus* ecotypes, predator ages, and changes to the microbial community of the prey field. Our results provide quantitative retention rates over these conditions that could fill gaps in our understanding of picocyanobacteria mortality, selective pressures, predictions of microbial biogeochemistry and oceanic carbon flux.

## Results

### Prochlorococcus retention rates increase with increasing prey concentration

Because increasing prey concentration leads to increased ingestion of eukaryotic by appendicularians (Selander & Tiselius, 2003), we addressed how *Prochlorococcus* concentration in the prey field influenced retention in the appendicularian. For both strains, MED4 (Figure 2A) and MIT9312 (Figure 2B), retention rates increased with increasing prey concentration. Retention rates ranged from tens of cells per minute per animal at the lowest concentrations to thousands of cells per minute per animal at the highest prey concentrations. Retention rates were not saturated at the highest *Prochlorococcus* concentrations, which are the highest concentrations relevant to *Prochlorococcus* in the ocean. Converting clearance rates of *Prochlorococcus* by appendicularians measured in natural ecosystems (Dadon-Pilosof et al., 2023; Scheinberg et al., 2005) to retention rates, showed that the rates we measured were comparable to the range of feeding rates wild appendicularians exert on natural communities of *Prochlorococcus* (Supplemental Figure 2).

**Figure 2.**
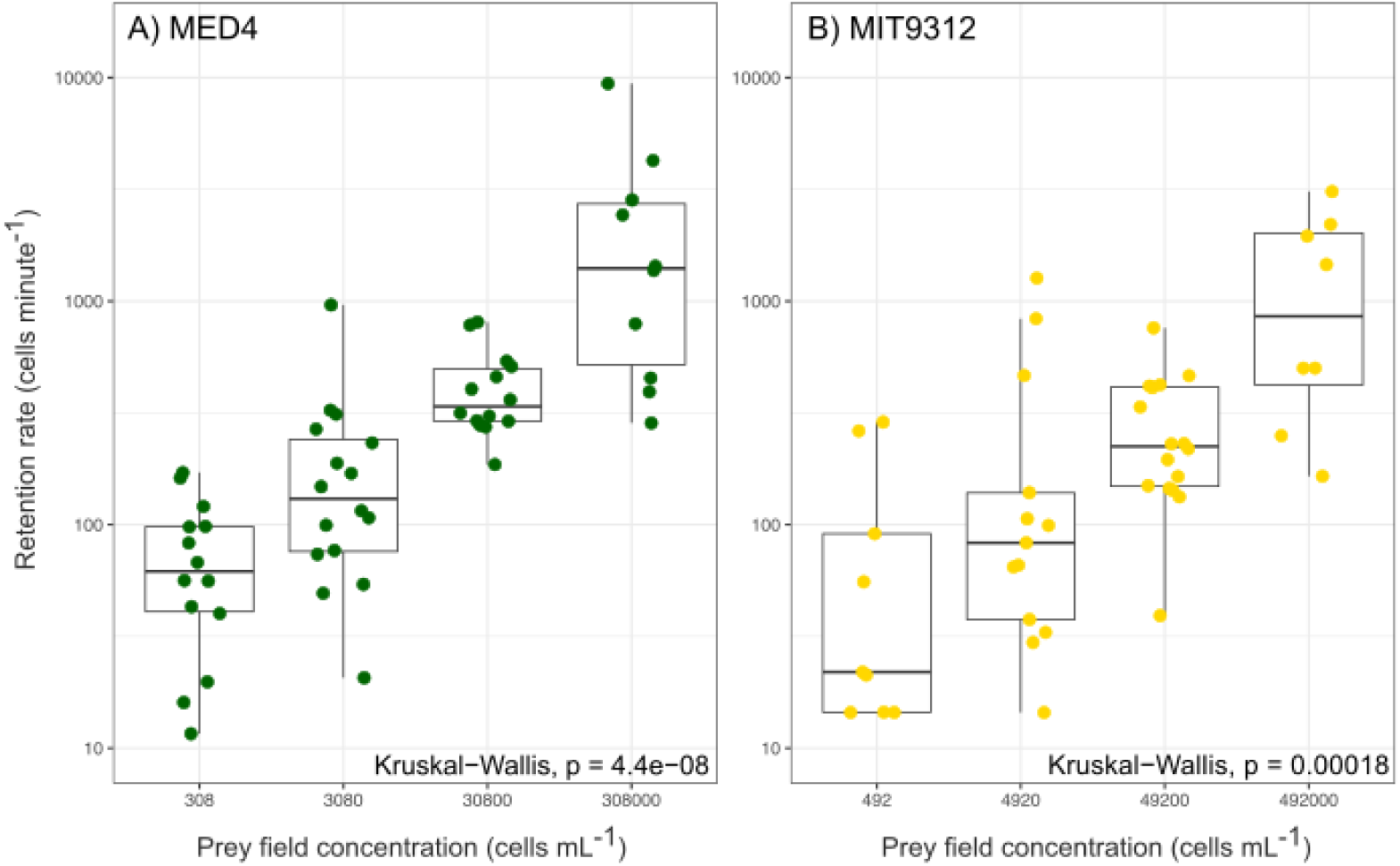
Retention rates of *Prochlorococcus* over ecologically relevant concentrations for A) MED4 and B) MIT9312.

### Appendicularians retain MED4 and MIT9312 at the same rates

Because *Prochlorococcus* is represented by diverse ecotypes, some of which dominate specific habitats and others that coexist (Johnson et al., 2006; Moore et al., 1998; Thompson et al., 2021), we tested whether strains from the two globally dominant ecotypes (HLI vs HLII) were differentially removed by appendicularians. This experiment was performed over a range of prey concentrations, because prey concentrations influence retention rate (Figure 2). We found that appendicularians grazed MED4 and MIT9312 at the same rate (*z-score* = 1.608, p-value > 0.05, Figure 3), despite slight differences between these two cell types in their diameter (MIT9312 is 0.83 µm and MED4 is 0.9 µm via Coulter Counter, (Ribalet et al., 2019)) and any unknown morphological or chemical differences that could result from their distinct genomes (Coleman et al., 2006; Kettler et al., 2007).

**Figure 3.**
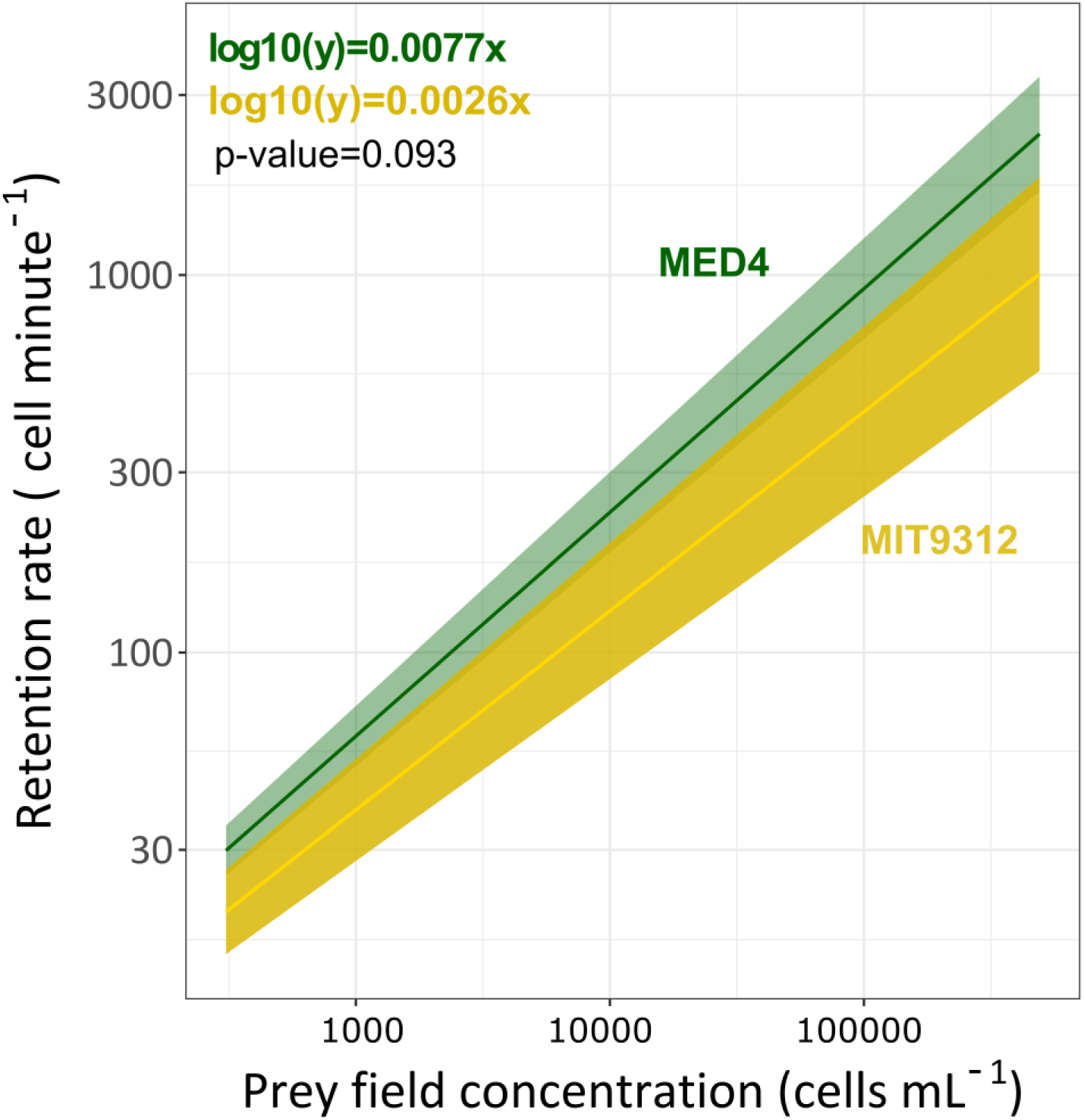
Comparison of the models of appendicularian retention rates vs *Prochlorococcus* concentration of two different *Prochlorococc*us strains (MED4 - green and MIT9312 - yellow).

### Advanced appendicularian ages retain more Prochlorococcus

Next we compared *Prochlorococcus* retention at different appendicularian ages, since mesh pore size (Deibel & Lee, 1992; Tiselius et al., 2003) and filtration rates (King et al., 1980; Lawrence et al., 2018) change during appendicularian development (Touratier et al., 2003). In each experiment, we compared two ages of appendicularians on the same prey field because no more than two ages were available each day at the culture facility. We did not compare experiments done on different days, due to slight differences in prey concentration and nutrient status each day, which would influence retention rates (Figure 2). Thus, only experiments done on the same day, with the same prey field preparation, were compared.

For both MIT9312 and MED4, retention rates were higher at more advanced ages (Figure 4A-C). For MED4, D5 retention rates were higher than D4 (Wilcoxon, p-value < 0.01; Figure 4A) and D7 retention rates were higher than D6 (Wilcoxon, p-value < 0.01, Figure 4B). For MIT9312, retention rate by D5 was higher than D4 (Wilcoxon, p-value < 0.01; Figure 4C). We did not test D6 and D7 appendicularians for MIT9312 due to methodological constraints, see Methods.

**Figure 4.**
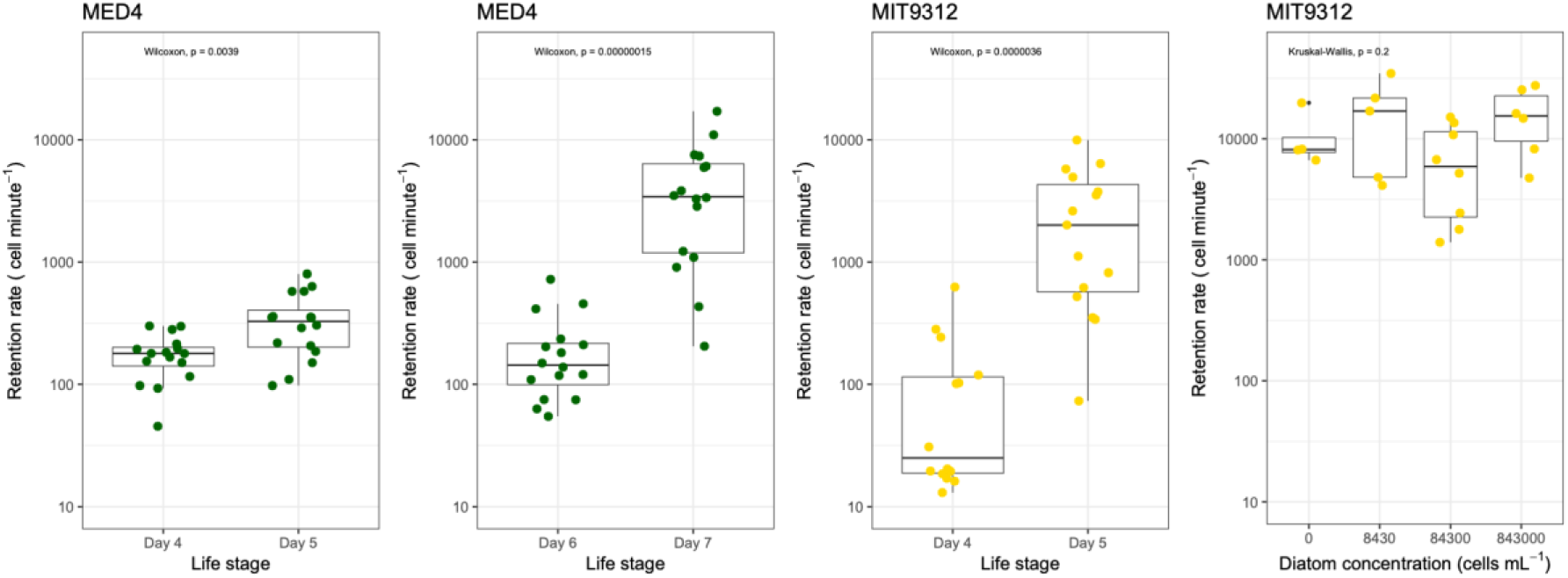
The effect of appendicularian ages (A-C) and the presence of other prey (i.e. Chaetocerous, D) on *Prochlorococcus* retention rates. *Prochlorococcus* strains MED4 (A+B) and MIT9312 (C+D) were used. Differences between treatments (x-axis) were tested with Kruskal-Wallis.

### Availability of eukaryotic phytoplankton prey did not change Prochlorococcus retention

We examined whether the presence of a large eukaryotic phytoplankton prey changed rates of *Prochlorococcus* retention. The rationale for this experiment was based on past observations that large and abundant prey can clog filtering meshes and alter appendicularian feeding behavior and filtration rates (Tiselius et al., 2003). We found no statistical support for differences in *Prochlorococcus* retention across four orders of magnitude difference in the abundance of the chain-forming diatom *Chaetocerous calcitrans* (Kruskal-Wallis, p-value > 0.05, Figure 4D).

## Discussion

This work examined the factors that influence the removal of *Prochlorococcus* from the ocean by *O. dioica*, a ubiquitous gelatinous filter feeder. While evidence exists for *Prochlorococcus* loss by appendicularians (Dadon-Pilosof et al., 2023; Scheinberg et al., 2005), this is the first study to address the factors that control the dynamics of the feeding interaction experimentally. Through feeding experiments with lab-reared appendicularians and cultivated *Prochlorococcus*, we demonstrated that increasing prey concentration and advancing predator age increased retention rates of *Prochlorococcus* while *Prochlorococcus* ecotype identity and the presence of larger prey cells did not influence retention. This quantitative insight will fill gaps in global ecosystem and carbon cycle models, many of which lack parameterization for filter-feeding predators of marine microorganisms such as appendicularians. Further, this work establishes a tractable experimental system that can be used to address questions on *Prochlorococcus* ecological and evolutionary responses to non-viral predation, an area that is almost completely unknown yet central to microbial ecology.

### Prochlorococcus loss rates by appendicularians vary across appendicularian and prey properties

Our work revealed that the removal of *Prochlorococcus* by appendicularians is dynamic due to both predator and prey characteristics. These findings extend previous work that established appendicularians as predators of *Prochlorococcus* (Dadon-Pilosof et al., 2023; Scheinberg et al., 2005), but due to working with wild populations, could not control predator and prey characteristics experimentally.

For the two globally dominant *Prochlorococcus* clades (HLI and HLII), appendicularian retention rates increased with increasing prey concentrations (Figures 2), which has important implications for representing this relationship in ecosystem models, such as those used to estimate the global microbial biogeography and carbon cycling (Dutkiewicz et al., 2024). For informing grazing rates in ecosystem models, we found that the highest known concentrations of *Prochlorococcus* in the ocean (up to 250,000 cells per mL (Flombaum et al., 2013)) showed no evidence of saturating retention rates (Figure 2, 3). While some maximum limit likely exists, it is unlikely for appendicularians to encounter high enough concentrations of *Prochlorococcus* to saturate feeding under current climate conditions. However, if *Prochlorococcus* concentrations increase with climate change (Bian et al., 2023; Flombaum et al., 2013), such high concentrations could become relevant.

Overall, this work suggests that the relationship between prey concentration and appendicularian feeding rate is different between smaller marine bacteria and larger eukaryotic phytoplankton, which can clog filters and slow appendicularian feeding at high concentrations or when defenses (i.e. spines) exist (Selander & Tiselius, 2003). This variation in maximum clearance rates by prey type underscores the need to quantify prey-specific retention rates to create a framework to accurately predict *in-situ* grazing rates on marine bacteria and archaea.

We also discovered that more advanced appendicularian ages consumed more *Prochlorococcus* (Figure 4A-C), which is consistent with grazing on eukaryotic phytoplankton (King et al., 1980; Lombard et al., 2009) and viruses (Lawrence et al., 2018). A likely mechanism is the increase of individual appendicularian size and filtration rate as ages advances (Lombard et al., 2009; Touratier et al., 2003). However, appendicularian growth rate over the life cycle transitions non-linearly from somatic growth to sexual maturation near spawning (i.e. at D7 under our conditions) (Troedsson et al., 2002). Thus, the relationship between appendicularian growth stage, growth rate, and prey retention remains unclear as the timing of the appendicularian cohorts we used limited us to pair-wise comparisons of ages (i.e. D4 vs D5, and D6 vs D7). Quantifying retention rate simultaneously across many ages, not just pairwise comparisons, would address the relationship between individual grazer size, age, filtration rate, and *Prochlorococcus* retention. Such investigation may also be relevant to predictions of the future ocean, as appendicularian fecundity could increase with ocean acidity (Taucher et al., 2024), and thus have more complex effects on their growth rates, nutrient requirements, and predation.

### Prochlorococcus retention rates are constant despite the presence of larger phytoplankton prey

We tested how the surrounding microbial community influenced *Prochlorococcus* retention because appendicularian filters are sensitive to clogging due to high prey biomass, specific prey features such as spines, and feeding efforts can decrease with increasing food concentrations (Selander & Tiselius, 2003; Tiselius et al., 2003) We found no statistical support for different retention rates of *Prochlorococcus* when *C.calcitrans* were present (Figure 4D). Thus, the presence of diatoms at concentration typical of the coastal ocean (10^5^ cells per mL), did not change feeding through mechanisms such as clogging on the inlet filter, food concentration filter, pharyngeal filter, or reduced feeding effort, all of which has been observed for larger prey species, but not smaller eukaryotic phytoplankton prey (Troedsson et al., 2007). The impact of larger prey (> 3 µm diameter) on *Prochlorococcus* grazing is like that for other small prey (∼0.5-1 µm). This finding suggests that the retention rates we have measured for appendicularian age (Figure 4), and prey concentration (Figure 2), would remain relevant under different microbial communities associated with blooms, upwelling, fronts, or large-scale ocean gradients.

### *Appendicularians graze equally on the most abundant* Prochlorococcus *ecotypes*

While *Prochlorococcus* viruses are host-specific (Sullivan et al., 2003), and protists feed selectively between ecotypes of the other major picocyanobacterium *Synechococcus* (Apple et al., 2011), it is unknown whether gelatinous zooplankton predators select between picocyanobacterial ecotypes. Determining whether non-viral predators choose between ecotypes is important to understand if predator interactions contribute to their distinct global biogeographic patterns or exert different evolutionary pressure on distinct ecotypes.

We found that representatives from the two most abundant *Prochlorococcus* clades (HLI and HLII) were retained equally by appendicularians (Figure 3), which suggests that the subtle differences between these two cell types do not influence capture by the appendicularians. While each ecotype contains hundreds of strain-specific genes (Coleman et al., 2006), this work suggests that they do not encode for any predation defenses (e.g. surface molecules) that are effective against appendicularians. MED4 and MIT9312 are only very slightly different in spherical diameter and carbon content (Ribalet et al., 2019), which also explains equal retention by appendicularians, as size and shape are important selective factors (Conley & Sutherland, 2017). However, there is still the possibility that appendicularians ingest these two cell types differently even after equal retention in the food concentrating filter. Following the food concentrating filter is the pharyngeal filter (Figure 1A), which presents another layer of selection before ingestion at the animals’ mouth (Deibel & Lee, 1992). Studies that quantify these cell types in the house separately from the animal, such as a recent study with artificial beads of different surface charge (Hiebert et al. In Review), could reveal if selection occurs at the point of ingestion rather than attachment to the mucous mesh.

One major difference between these two strains is their optimal latitudes, with MED4 dominant at temperate latitudes and MIT9312 at subtropical and tropical latitudes (Johnson et al., 2006). Appendicularians inhabit tropical, subtropical, and temperate waters and so share ranges with both ecotypes (Gorsky & Fenaux, 1998). The equal retention of MED4 and MIT9312 suggests that appendicularians could feed on *Prochlorococcus* over its entire global range. Thus, appendicularians may be a predator of *Prochlorococcus* on a global scale. Despite being globally abundant, appendicularians are underestimated due to their fragility and damage in nets, and are only just emerging as globally dominant predators through use of in situ imaging (Drago et al., 2022; Jaspers et al., 2023; Panaïotis et al., 2023; Schmid et al., 2023). Quantification of appendicularians in *Prochlorococcus* habitats would help estimate the global impact and dynamics of appendicularian feeding on *Prochlorococcus*.

### Unrecognized pathways for Prochlorococcus in marine carbon cycles and food webs revealed by appendicularian predation

Predation by appendicularians offers a new mechanism by which *Prochlorococcus* integrates into marine carbon cycles as appendicularian entry into the carbon cycle is well understood and multifaceted (Figure 5). Appendicularians contribute to carbon export through their sinking fecal pellets (Conley & Sutherland, 2017; Gorsky et al., 2005; Vargas et al., 2002) and routinely discarded houses (Jaspers et al., 2023). Both fecal pellets and the discarded houses can release remineralized carbon as well as intact cells as they sink (Lombard & Kiørboe, 2010). Appendicularian predation could explain observations of *Prochlorococcus* export carbon to the deep ocean and intermediate depths (De Martini et al., 2018) and intact *Prochlorococcus* in aphotic waters (Jiao et al., 2014). Indirectly, appendicularian predation on *Prochlorococcus* could alter contributions to the carbon cycle as *Prochlorococcus* produces key extracellular metabolites that fuel microbial metabolisms (Kujawinski et al., 2023). By removing *Prochlorococcus,* appendicularians may indirectly limit or change the delivery of these important carbon compounds to the microbial loop. Finally, appendicularians can actively reject prey at the pharyngeal filter (Lombard et al., 2011) after the cells have already been captured by interacting with sticky mucous fibers and feeding hydrodynamics (Hiebert et al., 2023). The fate and integrity of these rejected *Prochlorococcus* is uncertain, but interactions between cells and appendicularians may result in cell stress or compromised membrane integrity, resulting in increased susceptibility of *Prochlorococcus* to attack by other bacteria, protists, or viruses, and thus influence how *Prochlorococcus*-materials enter marine carbon cycles.

**Figure 5.**
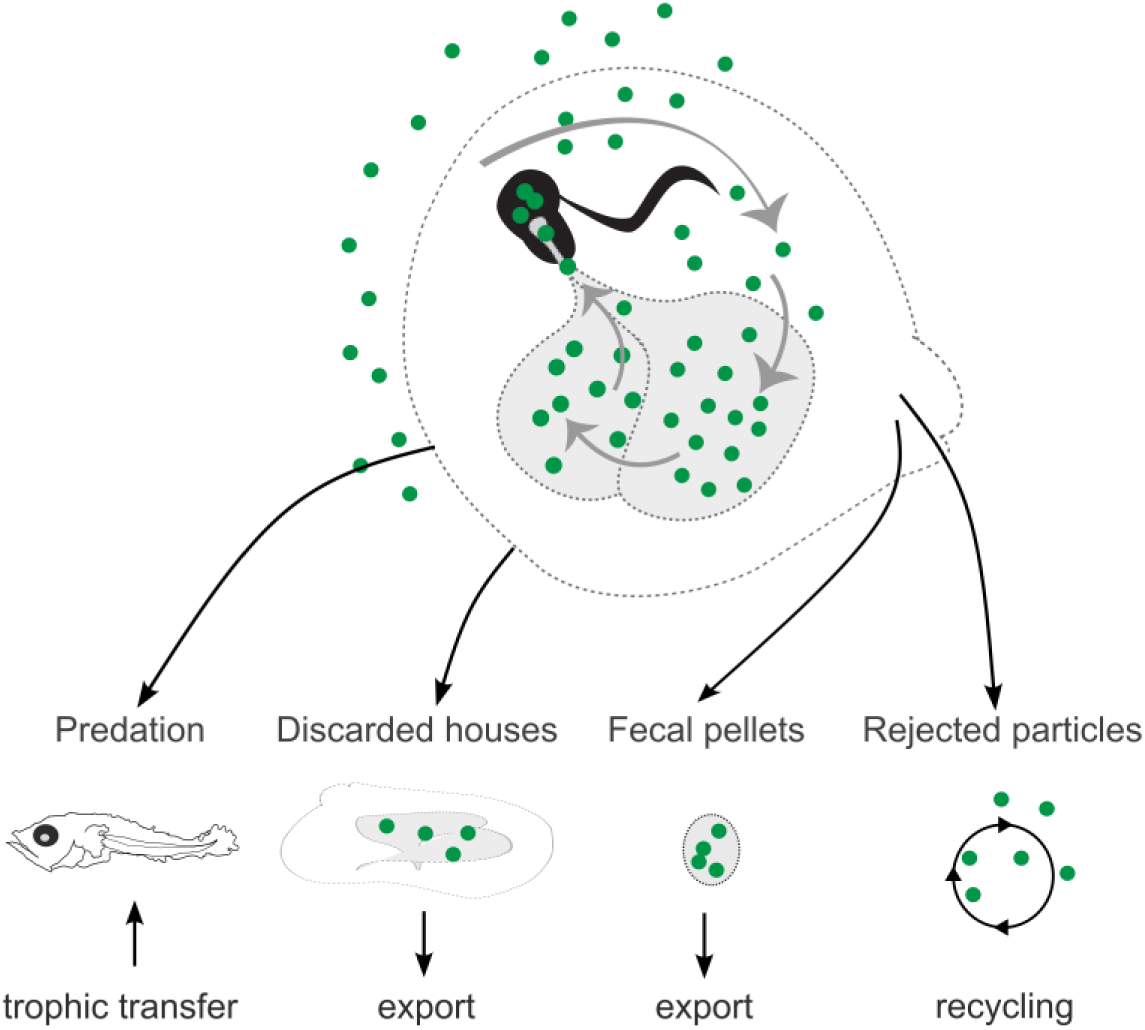
Diagram of mechanisms by which *Prochlorococcus* (green circles) could contribute to oceanic carbon cycles because of predation by appendicularians. Appendicularian features as labeled in Figure 1A. A) Ingestion and assimilation to support appendicularian growth followed by predation by larval fishes (e.g. tuna) and trophic transfer. B) Incorporation into discarded house, which becomes marine snow and contributes to carbon export. C) Ingestion and incorporation into fecal pellets, which sink and contribute to carbon export (black arrows). D) Rejection back into seawater to continue contributing towards carbon cycling in the microbial loop. Diagram modified from Hiebert et al. (In Review).

Appendicularian predation also offers a previously unrecognized role for *Prochlorococcus* in support of higher trophic levels (Figure 5). As an important and sometimes sole food source for several species of larval fishes (e.g. larval tuna) (Kalarus & Panasiuk, 2021; Llopiz et al., 2010), appendicularians create a one-step link between *Prochlorococcus* and high trophic levels in the subtropical oceans, where *Prochlorococcus* dominates. Feeding on *Prochlorococcus* and other marine bacteria, supports the idea that appendicularians, and other gelatinous zooplankton, can deviate from predator-prey size ratios in traditional marine food webs and form stable trophic links in changing ecosystems (Gorsky & Fenaux, 1998; Jaspers et al., 2023).

## Conclusion

Our work shows that appendicularians are dynamic predators of *Prochlorococcus.* Retention of *Prochlorococcus* increased with concentration and appendicularian age, suggesting that predation rates may change over latitudinal gradients in *Prochlorococcus* and through the progression of appendicularian blooms. Because retention rates did not vary between the dominant *Prochlorococcus* ecotypes, or when other phytoplankton were present, these relationships may be relevant to global-scale predictions of microbial mortality, biogeography, and carbon cycling. This work demonstrates that the unique feeding mechanism of appendicularians provides previously unrecognized carbon pathways for *Prochlorococcus*, such as a mechanism for export. This work highlights how further insight into microbial ecology can be reached with robust culture systems of predator and prey.

## Materials and Methods

### Appendicularian cultivation

The cultivation of appendicularians followed Bouquet et al., (2009) and was performed at the Michael Sars Centre in Bergen, Norway. Briefly, appendicularians (*O. diocia*) were reared in 8 L tanks filled with 6 L of filtered seawater (< 1 μm) (FSW) which was treated with activated charcoal and UV-light. Cultivation temperature was 12.2 - 13.0°C, which set the appendicularian life cycle progression to 7 days. The appendicularians were transferred to experimental beakers using wide bore pipettes and observed for tail beating prior to use in experiments.

### Prochlorococcus cultivation

*Prochlorococcus* strains MIT9312 and MED4 were acclimated to 20 °C, 12-hour light intervals at 25 μE m^-2^ s^-1^, and Artificial seawater (AMP1) with PRO99 medium amendments (Moore et al., 2007). *Prochlorococcus* cultures were transported to Bergen, Norway in cell culture flasks (Thermo Fisher Scientific Inc., Waltham, Massachusetts) then stored at room temperature (about 20 °C).

### Experimental designs

All experimental prey fields were made with filtered seawater amended with live *Prochlorococcus*. Details for each specific experiment are described below, and common elements are as follows: Prey fields were prepared in 1-L beakers, stirred to ensure even mixing, and then divided into duplicate beakers for each treatment. Using an assumed filtration rate of 12 mL hr^-1^ (Alldredge, 1981), the number of appendicularians added to each prey field was 8-12, few enough to prevent a significant change in the prey field over the course of the experiment, which was confirmed for each experiment (Supplemental Figure 1). For each experiment, swimming appendicularians with intact houses were transferred individually from their 8-liter culture beakers to the incubation chambers with experimental prey fields using a wide bore pipette. Experimental incubations took place in a sea table at 12 °C with a rotating plastic paddle to ensure even mixing throughout the experiment. After 10 minutes each appendicularian was individually transferred to a watch glass containing 2 mL of filtered seawater (along with 0.5-1 mL background prey field), and the time of removal was recorded. Each appendicularian was viewed under a compound microscope to ensure that the house was still attached, then fixed individually in 100 μL of DNA/RNA Shield (Zymo Research Corporation, Irvine, California), stored at room temperature for transport to the home lab, then at -80 °C until DNA extraction.

Experiment-specific treatments are as follows: For experiments testing appendicularian feeding across different *Prochlorococcus* concentrations, D5 appendicularians were used. For each prey concentration, two replicate chambers were used. For experiments testing retention rates of *Prochlorococcus* by the appendicularians across different ages (D1, D2, D4, and D5) 8.8×10^4^ MIT9312 *Prochlorococcus* cells mL^-1^ were fed to the D1 and D2 appendicularians and 8.5×10^3^ MIT9312 *Prochlorococcus* cells mL^-1^ were fed to the D4 and D5 appendicularians. For experiments testing retention rates of *Prochlorococcus* in the presence of different concentrations of a larger prey, *Prochlorococcus* concentrations were constant in each treatment and Chaetocerous concentrations included: none, ∼8400 cells mL^-1^, ∼84000 cells mL^-1^, and ∼840000 cells mL^-1,^ to replicate naturally occurring concentrations of large eukaryotic phytoplankton in the ocean.

To make sure the *Prochlorococcus* prey field did not change during the experiments, all prey fields were sampled prior to (t_0_) and at the end of the experiment (t_F_), in triplicate, and assessed for *Prochlorococcus* concentration by qPCR (see details below). There were no statistically supported differences in the prey fields before and after the experiments (Supplemental Figure 1).

### DNA extraction and Quantitative PCR

DNA was extracted from appendicularians (house and animal sampled together) using the Quick-DNA™ Miniprep Kit (Zymo Research Corporation, Irvine, California) with modifications including using two 400 μL aliquots of lysate during the extraction process. Extracted DNA concentrations were quantified using the NanoDrop™ One Microvolume UV-Vis Spectrophotometer (Thermo Fisher Scientific Inc., Waltham, Massachusetts).

Quantitative PCR to detect *Prochlorococcus* in the prey field and in each appendicularian was performed using the Applied Biosystems® ViiA™ 7 Real-Time PCR System with the Power SYBR™ Green PCR Master Mix (Applied Biosystems, Waltham, MA), following published *Prochlorococcus* eMED4 and eMIT9312 assay protocols (Zinser et al., 2006). gBlocks™ Gene Fragments (Integrated DNA Technologies, Coralville, IA) serial dilutions (10^0^ to 10^6^ gene copies per reaction) were used for the standards and were diluted using DNase/RNase Free Water (Zymo Research Corporation, Irvine, California) (Supplemental Table 1). Standards were run in triplicate. No-template controls with water were run in duplicate. Melt curve analysis was performed for each qPCR run and confirmed the presence of a single amplicon.

To calculate *Prochlorococcus* gene copies per appendicularians, the mean copy number from each triplicate qPCR reaction was multiplied by the volume of elution and divided by percent of sample (84%) transferred to the spin column following the lysate step of DNA extraction. The total gene copy count (i.e. *Prochlorococcus* cell count) per appendicularian sample collected was divided by the incubation time for each appendicularian to yield individuals’ grazing rates as cells minute^-1^. For prey field samples, the total cell count per sample was divided by the volume of water collected, adjusted as described above, to yield the prey field concentrations expressed as cells mL^-1^.

To estimate the carbon uptake through grazing on *Prochlorococcus* across the range of concentrations, the retention rate in minutes (cells appendicularian^-1^ minute^-1^) was converted to an hourly retention rate (cells appendicularian^-1^ hour^-1^). This hourly rate was multiplied by 24 hours to determine the daily retention rate (cells appendicularian^-1^ day^-1^). The daily retention rate was multiplied by the average carbon content of a *Prochlorococcus* cell (46 femtograms cell^-1^) (Bertilsson et al., 2003), resulting in a daily carbon uptake from *Prochlorococcus* per individual appendicularian.

### Data analysis and visualization

All data analysis and visualization were performed with R version 4.3.2 in RStudio (Posit team, 2024; R Core Team, 2023). Plots were made using *ggplot2* (Wickham et al., 2016), *dplyr* (Wickham et al., 2023), *broom* (Robinson et al., 2024), *ggpubr* (Kassambara, 2023), and *ggtext* (Wilke & Wiernik, Brenton M., 2022). The values for variation and error were calculated using the Wilcoxon t-test for the replicate samples. The Kruskal-Wallis test was utilized to compare retention rates (copy number individual^-1^ time of incubation^-1^) across multiple groups.

## Author contributions

Description of author contributions follows the “CRediT” taxonomy (Brand et al., 2015). Conceptualization: A.W.T, C.P.S., T.H., K.R.S.; Methodology: A.W.T., C.P.S, A.H., A.U.K., A.A., D.C.; Formal Analysis: A.W.T., C.P.S, A.H., A.U.K.; Investigation: A.W.T., C.P.S, A.H., A.U.K., K.R.S.; Writing: A.W.T., C.P.S, A.H., A.U.K., K.R.S., T.H.; Visualization: A.W.T., C.P.S.; Supervision: A.W.T.; Funding Acquisition: A.W.T. and K.R.S.

## Acknowledgements

Funding was provided by Simons Foundation award LS-ECIAMEE-00001481 to AWT and NSF OCE 1851537 to KRS. We are grateful to the Molecular Testing Labs (Vancouver, WA), including Charles Sailey, Jill Whitford, and Andrew Chastain, for the donation and time spent optimizing the ViiA7 qPCR machine. Thank you to Kylee Brevick for qPCR support and calibration.

